# Neuropil aggrecan, not perineuronal nets, closes the critical period for visual plasticity

**DOI:** 10.64898/2026.07.26.740801

**Authors:** Emily C. Crouse, Thomas C. Brown, Xiaokuang Ma, Hongtao Ma, Kylee R. Karpowich, Kieran Barber, Peng Chen, Shenfeng Qiu, Aaron W. McGee

## Abstract

Perineuronal nets (PNNs) are widely reported to close developmental critical periods and restrict experience-dependent plasticity. Here we tested this model by selectively eliminating gene expression for aggrecan (*Acan*), an essential component of PNNs. In visual cortex, PNNs predominantly ensheath parvalbumin-positive (PV+) interneurons. Deletion of *Acan* in inhibitory neurons eliminated PNNs but did not prevent closure of the critical period. By comparison, deletion of *Acan* in all neurons, or only in excitatory forebrain neurons, sustained critical-period plasticity in adult mice. Visual plasticity in adults was associated with reduced cortical excitatory synaptic inputs onto layer 2/3 PV+ interneurons and increased expression of some immediate early genes in visual cortex but normal response strength and tuning properties of layer 2/3 excitatory neurons. These findings rectify long-standing models that misattribute closure of the critical period to PNNs and identify that aggrecan expressed by excitatory neurons resident in the surrounding neuropil limits visual plasticity.

## Introduction

Perineuronal nets (PNNs) were first described by Camillio Golgi in the 1890s from sections of spinal cord stained with his method; he proposed they provided protection and structural support in the adult nervous system (*1*). In contrast, Santiago Ramon y Cajal asserted that PNNs were an artefact of tissue fixation (*2*). Despite advances in understanding the molecular composition of PNNs (*3*), direct evidence for the principal functions of these extracellular structures has proven elusive for more than a century.

PNNs are a form of condensed extracellular matrix that contains members of the lectican family of chondroitin sulfate proteoglycans (CSPGs), tenascin glycoproteins, and hyaluronan (*3*, *4*). The proteoglycan link protein 1 (Hapln1) is also associated with PNNs in cerebral cortex and tethers CSGPs to the meshwork of hyaluronan (*4*, *5*). CSPGs are decorated with long chains of chondroitin sulfate glycosaminoglycans (CS-GAGs) and have molecular weights that are often hundreds of kilodaltons because of these post-translational modifications (*6*). Consequently, CSPGs have limited compatibility with common molecular biological approaches for evaluating function such as immunoprecipitation and immunoblotting.

Aggrecan is the principal CSPG in PNNs but is often accompanied by neurocan, versican, and brevican (*7*). Mice with a mutation in *Acan*, the gene encoding aggrecan, were first described as the *cartilage matrix deficiency* (*cmd*) mouse; homozygous mutants die at birth from respiratory failure (*8*, *9*). By comparison, ‘knock-out’ mutants for neurocan, brevican, and tenascin-R, as well as a quadruple mutant with tenascin-C, display only a partial reduction of the number PNNs (*10*–*14*). *Hapln1* mutant mice also have attenuated PNNs despite normal expression levels of CSPGs (*5*).

Several lines of indirect evidence implicate PNNs in closing the critical period for visual plasticity. PNNs are present in primary visual cortex (V1) of rodents during the critical period, but their abundance and staining intensity with wisteria floribunda agglutinin (WFA) increase across the closure of the critical period and plateau later in adulthood (*15*, *16*). In a landmark study, Pizzorusso et al. (2002) injected the enzyme chondroitinase ABC (chABC) into V1 of adult rats (*15*). ChABC degrades not only CS-GAGs, but also dermatan sulfates and hyaluronan (*17*, *18*). This treatment partially restored ocular dominance (OD) plasticity as measured with multi-unit electrophysiologic recordings (*15*). Similarly, injecting mouse V1 with an *AAV-Nestin-Cre* to delete a conditional ‘floxed’ allele (*flx*) of *Acan* in neurons and some glia abolished PNNs as reported by the loss of staining for WFA as well as the loss of immunostaining for aggrecan, neurocan, versican, brevican, phosphacan, tenascins, and Hapln1 (*19*). This expression pattern of Cre recombinase in adult *Acan flx/flx* mice also restored OD plasticity as measured with optical imaging of intrinsic signals (*19*). Although CSPGs are present both in PNNs and throughout the surrounding neuropil, the interpretation of these findings has been that the assembly of CSPGs into PNNs is required to restrict OD plasticity to the critical period (*4*, *7*, *15*, *19*). Here we performed a genetic dissection of the expression requirements for *Acan* to form PNNs and to close the critical period as a pathway to understanding the mechanisms that govern plasticity in the adult brain.

## Results

### PNNs do not close the critical period

As a prerequisite to dissecting the role of PNNs in closing the critical period for visual plasticity, we sought to reproduce the original finding that treatment of rodent V1 with chABC restores OD plasticity (*15*). Rather than repeatedly injecting purified enzyme into V1 as performed by Pizzorusso et al., we transduced cells in visual cortex with a combination of adeno-associated viruses (AAVs), *AAV-CMV-Pl-Cre-rBG* and the Cre-dependent *AAV-hSyn-DIO-ChABC-mCherry*, to drive the constitutive expression of chABC by neurons (*20*). Transduction with this combination of viruses abolished PNNs throughout V1 in the injected hemisphere but not the adjoining hemisphere (Fig. 1a and Fig. S1). PNNs were unaffected by cortical injection of the Cre-dependent *AAV-hSyn-DIO-ChABC-mCherry* alone (Fig. 1a). Mice expressing chABC in V1 were otherwise unremarkable in our experiments employing multi-unit electrophysiologic recordings and displayed typical visual responses, including high contralateral bias index (CBI) scores (Fig. 1b) (*21*, *22*).

**Figure 1.**
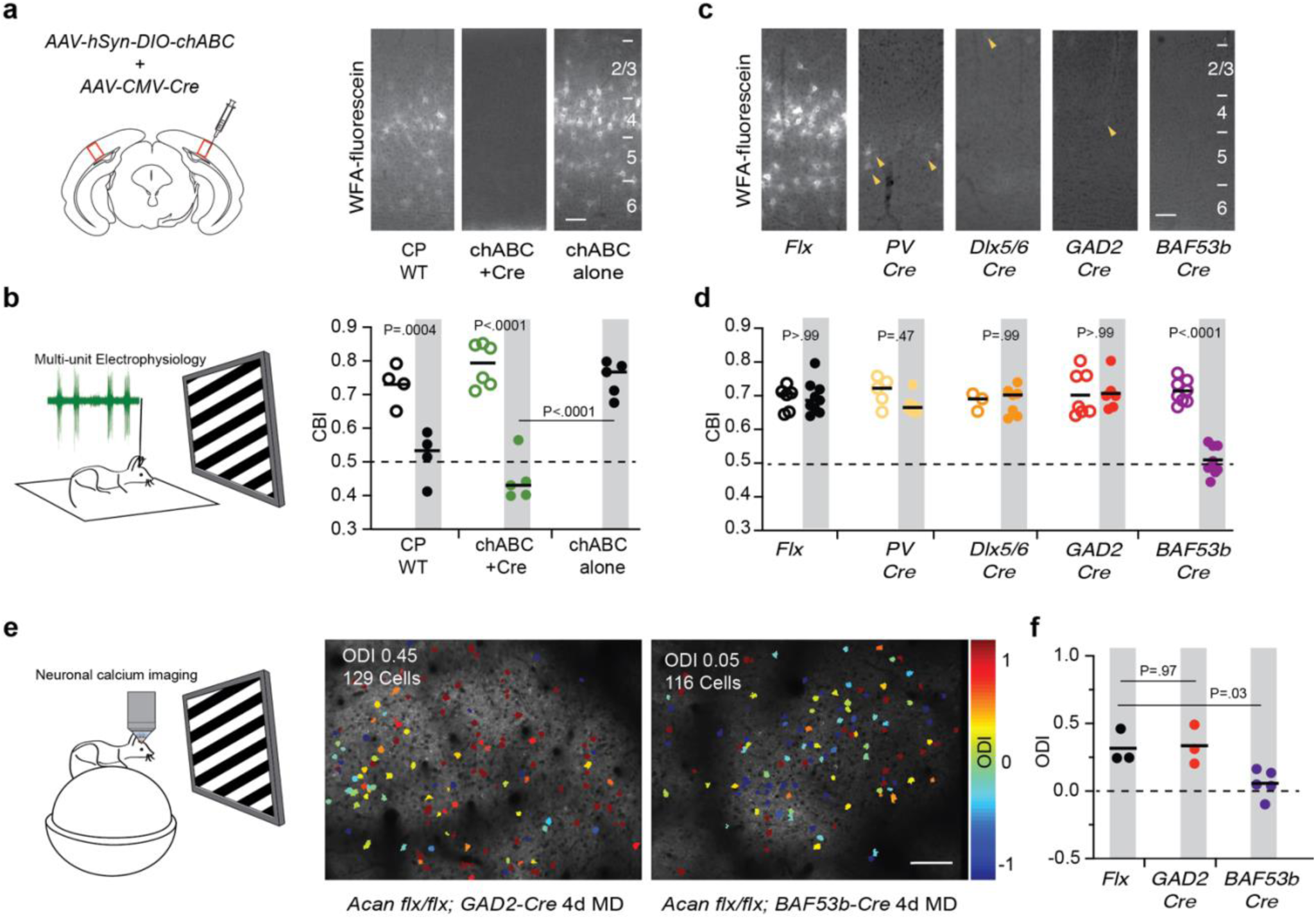
ChABC re-activates OD plasticity in adult mice which is phenocopied by loss of *Acan* in all neurons whereas eliminating *Acan* in inhibitory neurons abolishes PNNs but does not affect the closure of the critical period. **(a)** (*left*) Cartoon of a coronal section of mouse brain and injection sites for two AAVs (AAV-hSyn-DIO-chABC and AAV-CMV-Cre) to drive expression of chABC. (*right*) Coronal sections of V1 stained with WFA-fluorescein to label PNNs for a juvenile mouse (CP WT), the injected hemispheres for both chABC+Cre and the chABC alone negative control. Positions of cortical layers are indicated at right. Scale bar = 100 microns. **(b)** (*left*) Cartoon of the multi-unit recording method for anesthetized mice. (*right*) Mean Contralateral Bias Index (CBI) scores for untreated juvenile mice (CP WT; n=4/4), adult WT mice transduced with AAVs to express chABC (ChABC+Cre; n=6/5), adult WT mice transduced with the AAV for chABC alone that is inert without co-expression of Cre recombinase (ChABC alone; n=5). Open circles represent non-deprived mice. Closed circles and grey shading represent mice receiving 4-5d of MD prior to electrophysiologic recordings. The dashed line at 0.5 represents equal responsiveness to each eye. One-way ANOVA with Sidak correction for multiple comparisons (1-ANOVA). **(c)** Coronal sections of V1 stained with WFA-fluorescein to label PNNs in adult (P60-90) *Acan* flx/flx mice alone, or in combination with the indicated Cre driver. Yellow arrowheads indicate remaining PNNs. Positions of cortical layers are indicated at right. Scale bar = 100 microns. **(d)** Mean CBI scores as in panel (a) for adult *Acan flx/flx* mice alone (Flx; n = 6/9), or in combination with *PV-Cre* (n=5/8), *Dlx5/6-Cre* (n=3/7), *GAD2-Cre* (n=7/6), or *BAF53b-Cre* (n=8/9). **(e)** (*left*) Cartoon of the neuronal calcium imaging method for awake mice head-fixed on a spherical treadmill, and (*right*) example calcium imaging fields in V1 for *Acan flx/flx*; *GAD-Cre* or *BAF53b-Cre* mice after 4 days of MD. Colors correspond to Ocular Dominance Index (ODI) score where 1 (red) represents predominantly contralateral-responsive neurons, -1 (blue) represents predominantly ipsilateral-responsive neurons and the remaining color spectrum represents binocular-responsive neurons. **(f)** Mean ODI scores for adult *Flx* mice (= 3), *GAD2-Cre* (n=3) and *BAF53b-Cre* (n=5) following 4 days of MD from calcium imaging experiments. The dashed line at 0.0 represents equal responsiveness to each eye. 1-ANOVA.

Brief (4-5 days (d)) monocular deprivation (MD) is sufficient to yield saturating OD plasticity in juvenile mice during the developmental critical period (∼postnatal day (P) 21-32)(*21*, *23*, *24*). Expression of chABC restored the sensitivity of adult mice to brief MD. Injected adult mice that received MD prior to recording displayed lower CBI scores and shifted OD histograms that were similar to those of juvenile mice receiving MD during the critical period (Fig. 1b and Fig. S1). This OD plasticity was a consequence of expression of chABC rather than the injection procedure as transduction of cells in V1 with the inert *AAV-hSyn-DIO-ChABC-mCherry* alone yielded CBI scores typical for non-deprived mice that were significantly different from mice expressing chABC (P<.0001 for chABC+Cre 4d MD vs non-deprived or vs 4d MD chABC alone) (Fig. 1b). These experiments confirmed that degradation of sugar chains from CSPGs and other components in the extracellular matrix by chABC was indeed effective for re-activating OD plasticity (*15*).

Next, we dissected the requirement for *Acan* to close the critical period. Expression of *Acan* is relatively abundant in parvalbumin-positive (PV+) interneurons (*25*–*27*). Therefore, we examined the consequences of abolishing expression of *Acan* in PV+ interneurons with 3 distinct Cre drivers that provided overlapping expression patterns: *PV-Cre*, *Dlx5/6a-Cre*, and *GAD2-Cre*. *PV-Cre* is selective for cells expressing parvalbumin including PV+ interneurons (*28*). *Dlx5/6a-Cre* directs Cre recombinase expression to differentiating and migrating forebrain interneurons as early as embryonic day (E) 13.5, including PV+ interneurons (*29*, *30*). *GAD2-Cre* is selective for GABAergic interneurons, including PV+ interneurons (*31*). PNNs were numerous in coronal sections of V1 from adult *Acan flx/flx* control mice but were largely absent in sections of V1 from *Acan flx/flx; PV-Cre*, *Acan flx/flx; Dlx5/6a-Cre*, and *Acan flx/flx; GAD2-Cre* mice (Fig. 1c, Fig. S1). Deletion of *Acan* with any of these 3 different Cre drivers decimated the number of PNNs in V1.

We measured the sensitivity to MD of adult *Acan flx/flx* mice alone and in combination with each Cre driver selective for inhibitory neurons. None displayed OD plasticity (Fig. 1d and Fig. S1). The CBI scores of adult non-deprived *Acan flx/flx* mice were similar to adult wild-type (WT) mice and were not altered by 4-5d of MD (P = .99, non-deprived vs. 4d MD). *Acan flx/flx* mice expressing Cre recombinase in inhibitory neurons lacked PNNs but did not display OD plasticity following 4-5d of MD (P = .47- >.99 for each genotype non-deprived vs. 4d MD). These findings reveal that PNNs do not close the critical period and contradict the conclusions of prior work asserting that these extracellular structures close the critical period and function as a brake on visual plasticity (*4*, *7*).

Given that expression of chABC in V1 restored OD plasticity but elimination of PNNs did not, we investigated whether broader deletion of *Acan*, akin to the deletion from *AAV-Nestin-Cre*, would prevent the closure of the critical period. We eliminated neuronal expression of *Acan* with *BAF53b-Cre*, a pan-neuronal Cre driver (*32*). *Acan flx/flx; BAF53b-Cre* mice also lacked PNNs (Fig. 1c). However, in contrast to *Acan flx/flx* mice in combination with Cre drivers selective for inhibitory neurons, deletion of *Acan* in all neurons with *BAF53b-Cre* sustained OD plasticity in adult mice following MD (P < .0001, non-deprived vs. 4d MD) (Fig. 1d and Fig. S1).

To confirm these results, we measured OD plasticity with calcium imaging at neuronal resolution in awake mice (Fig. 1e,f). We generated *Acan flx/flx* mice that possessed 2 transgenes, *Tg-tetO-GCaMP6s* and *Tg-Camk2a-tTA,* that together drive strong expression of the genetically-encoded calcium sensor GCAMP6s in excitatory neurons of the forebrain (*33*). Consistent with our previous calcium imaging experiments with adult WT mice (*34*), adult *Acan flx/flx* mice did not exhibit OD plasticity following 4-5d of MD and their Ocular Dominance Index (ODI) scores centered around 0.3, a value typical for non-deprived mice (Fig. 1f)(*34*–*36*). *Acan flx/flx; GAD2-Cre* mice also did not display OD plasticity as measured with calcium imaging of neuronal activity. By comparison, *Acan flx/flx; BAF53b-Cre* mice imaged after 4d of MD displayed OD plasticity and possessed lower ODI scores (Fig. 1f). These ODI scores were centered near 0.0 because of a relative reduction in the fraction neurons predominantly responsive to visual stimuli presented to the contralateral eye and a corresponding increase in the fraction of neurons responsive predominantly to the ipsilateral eye (Fig. S1) (*35*). OD plasticity measured with calcium imaging mirrored the OD plasticity we observed with multi-unit electrophysiologic recordings. Thus, we conclude that deletion of *Acan* in excitatory neurons is necessary to prevent the critical period from closing.

### OD plasticity is associated with reduced cortical inhibitory drive

Then, we examined if differences in intracortical synaptic circuitry contribute to the developmental visual plasticity displayed by adult *Acan flx/flx; BAF53b-Cre* mice. We again focused on PV+ interneurons as these cells display the highest neuronal expression of *Acan* (*25*, *27*). Cortical injection of chABC into adult mice increases the number and strength of excitatory synapses onto PV+ interneurons in V1 as reported by the frequency and amplitude of spontaneous excitatory postsynaptic currents (sEPSCs) (*37*). We labeled inhibitory neurons in V1 of adult mice with green fluorescent protein (GFP) by transducing neurons with *AAV-mDLX-GFP* for *Acan flx/flx* mice alone and in combination with either *GAD2-Cre* or *BAF53b-Cre* (Fig. S2) (*38*). Then we performed whole-cell recordings and laser scanning photostimulation (LSPS) to map the cortical excitatory synaptic inputs onto L2/3 PV+ interneurons in V1 (Fig. 2a-c). We targeted PV+ interneurons by their GFP fluorescence and intrinsic properties including high firing rates in response to current injection (Fig. 2d, Fig. S2). We confirmed expression of PV in recorded neurons with post-hoc immunostaining (Fig. S2). Elimination of PNNs did not yield detectable differences in the distribution or strength of excitatory synaptic inputs onto PV+ interneurons. The heatmaps and corresponding quantification of evoked EPSCs were similar between *Acan flx/flx* mice and *Acan flx/flx; GAD2-Cre* mice. By comparison, there was a moderate yet significant reduction in the strength of excitatory inputs onto PV+ interneurons in *Acan flx/flx; BAF53b-Cre* mice relative to *Acan flx/flx* controls (Fig. 2e-g).

**Figure 2.**
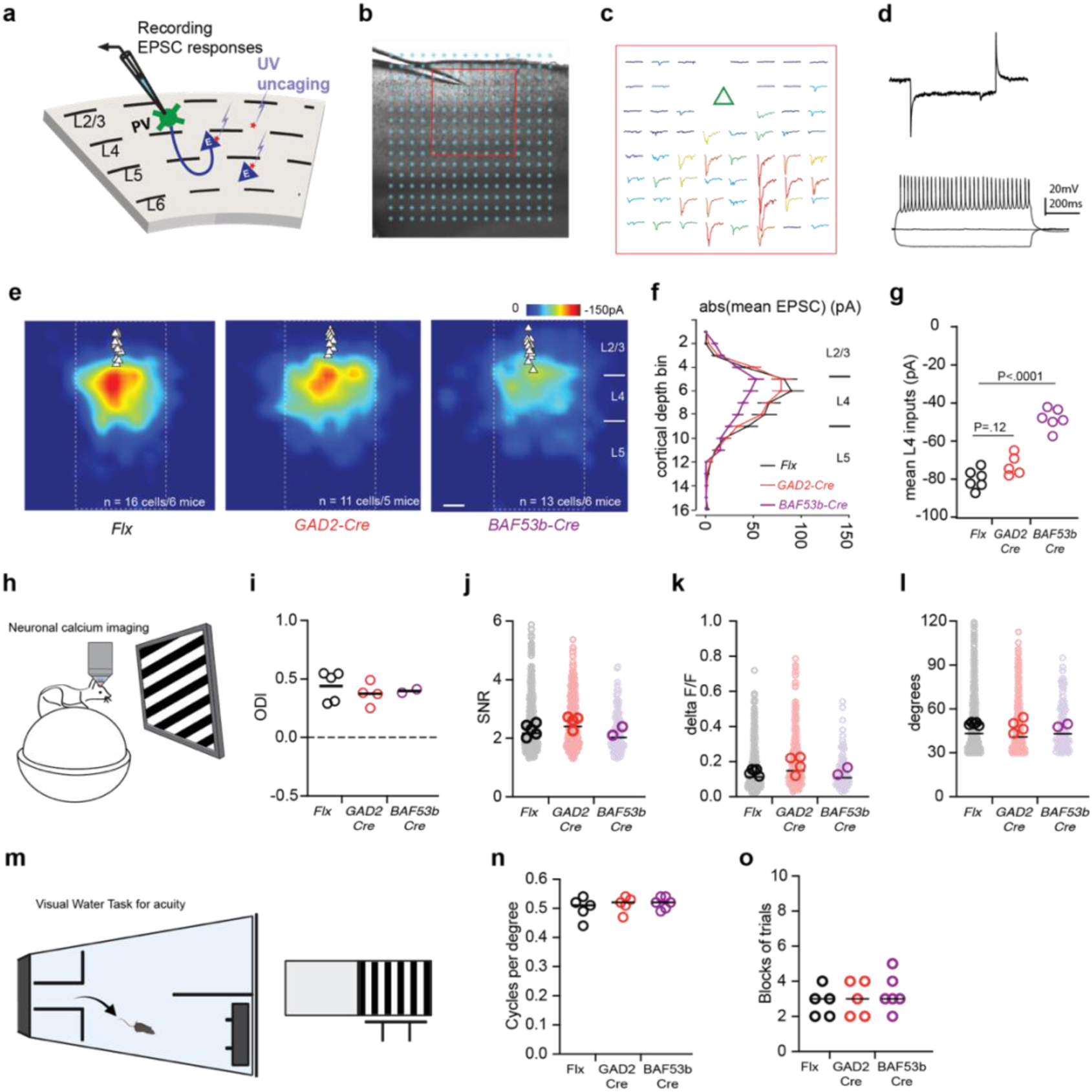
Sustained OD plasticity in adult *Acan flx/flx; BAF53b-Cre* mice is associated with reduced excitatory synaptic drive onto PV interneurons that is not mediated by loss of PNNs and does not alter tuning properties of cortical excitatory neurons or visual acuity. **(a)** Cartoon of the whole-cell recording from PV interneurons and LSPS setup. **(b)** Example whole-cell recording from a GFP-positive neurons overlaid with the 16X16 grid for glutamate uncaging. **(c)** Example responses following LSPS glutamate uncaging color-coded by amplitude at the central 8X8 stimulation grid locations surrounding the neuronal soma (indicated by red square in panel b) **(d)** (*top*) Current response of the L2/3 PV+ interneuron in response to a voltage pulse command. (*bottom*) Voltage responses to -50, 0, and 50pA current injections, which reveal the characteristic fast spiking of PV interneurons. **(e)** Heat maps of the aggregate distribution and strength of excitatory synaptic inputs (pA) onto PV+ interneurons. Triangles indicate soma location (*Flx*, n = 16 cells/6 mice; *GAD2-Cre*, 11 cells/5 mice; *BAF53b-Cre*, 13 cells/6 mice). Layer positions are indicated at right. Scale bar = 100 microns. **(f)** Mean absolute value (abs) of LSPS-evoked EPSC amplitudes from neurons in panel e binned by cortical depth of the stimulation site. **(g)** Mean input strengths per mouse for L4 inputs binned from the central 8 stimulation columns for each genotype. **(h)** Cartoon of the neuronal calcium imaging method for awake mice head-fixed on a spherical treadmill. **(i)** Mean ODI scores per mouse for non-deprived adult *Flx* mice (n=5), *GAD2-Cre* (n=4) and *BAF53b-Cre* (n=2) from calcium imaging experiments. The dashed line at 0.0 represents equal responsiveness to each eye. **(j)** SNR for all visually-evoked neurons responsive to the contralateral eye and the mean per mouse for non-deprived *Flx* (n=576 neurons, 5 mice), *GAD2-Cre* (n=356 neurons, 4 mice) and *BAF53b-Cre* (n=108 neurons, 2 mice). The horizonal lines represent the median for the neuronal responses for panels e-g. **(k)** The mean delta F/F per neuron and mouse for the same mice as in panel e. **(l)** The full width half-maximum of the orientation tuning. **(m)** Cartoon of the behavioral assay for measuring visual acuity. **(n)** Visual acuity for adult mice of each genotype: *Flx* (n=5), *GAD2-Cre* (n=5) and *BAF53b-Cre* (n=6). The horizonal lines represent the median. **(o)** The number of blocks to reach the criteria for acuity testing. One block = ten trials.

To determine if this reduction to intracortical excitatory synaptic drive onto PV+ interneurons in *Acan flx/flx; BAF53b-Cre* mice that was associated with sustained developmental visual plasticity in adult mice was also sufficient to alter the visually-evoked activity in V1, or the tuning properties of pyramidal neurons in L2/3, we performed *in vivo* calcium imaging on non-deprived mice (Fig. 2h-l). Non-deprived mice for each genotype displayed normal contralateral bias of visual responsiveness (Fig. 2i). We calculated the tuning properties of excitatory cortical neurons in adult *Acan flx/flx* mice alone or in combination with *GAD2-Cre* or *BAF53b-Cre* in response to visual stimuli presented to the contralateral eye (Fig. 2i-l). Visual responsiveness, as measured by both the signal to noise ratio (SNR) and the average normalized change in fluorescence (delta F/F), was similar for *Acan flx/flx* and *Acan flx/flx; BAF53b-Cre* mice (Fig. 2j,k). Additionally, the half-maximum full widths of the orientation tuning were also similar between genotypes (Fig. 2l). Consistent with these normal neuronal tuning properties, adult *Acan flx/flx* mice, *Acan flx/flx*; *GAD2-Cre* mice, and *Acan flx/flx*; *BAF53b-Cre* mice, possessed normal visual acuity as measured with the visual water task and required a similar number of blocks of trials to learn the task (Fig. 2m-o).

### Aggrecan expression in the neuropil closes the critical period

To assess whether deletion of *Acan* in inhibitory neurons of adult mice might yield a result more similar to treatment with chABC (*39*), we evaluated OD plasticity in adult *Acan flx/flx* mice that received injections into V1 of *AAV-hDLX-minBG-iCre-4X2C-WPRE3-BGHpA*, an AAV with a promoter that restricts Cre expression to inhibitory neurons(*40*) (Fig. S2). AAV injection into V1 abolished PNNs by 3 weeks post-injection while PNNs appeared normal in V1 of the adjoining hemisphere (Fig. 3a). Abolishing PNNs by deleting *Acan* in inhibitory neurons of adult *flx/flx* mice also did not re-activate OD plasticity as 4-5d of MD contralateral to the treated hemisphere yielded CBI scores similar to *Acan flx/flx* controls (Fig. 3b) (*41*).

**Figure 3.**
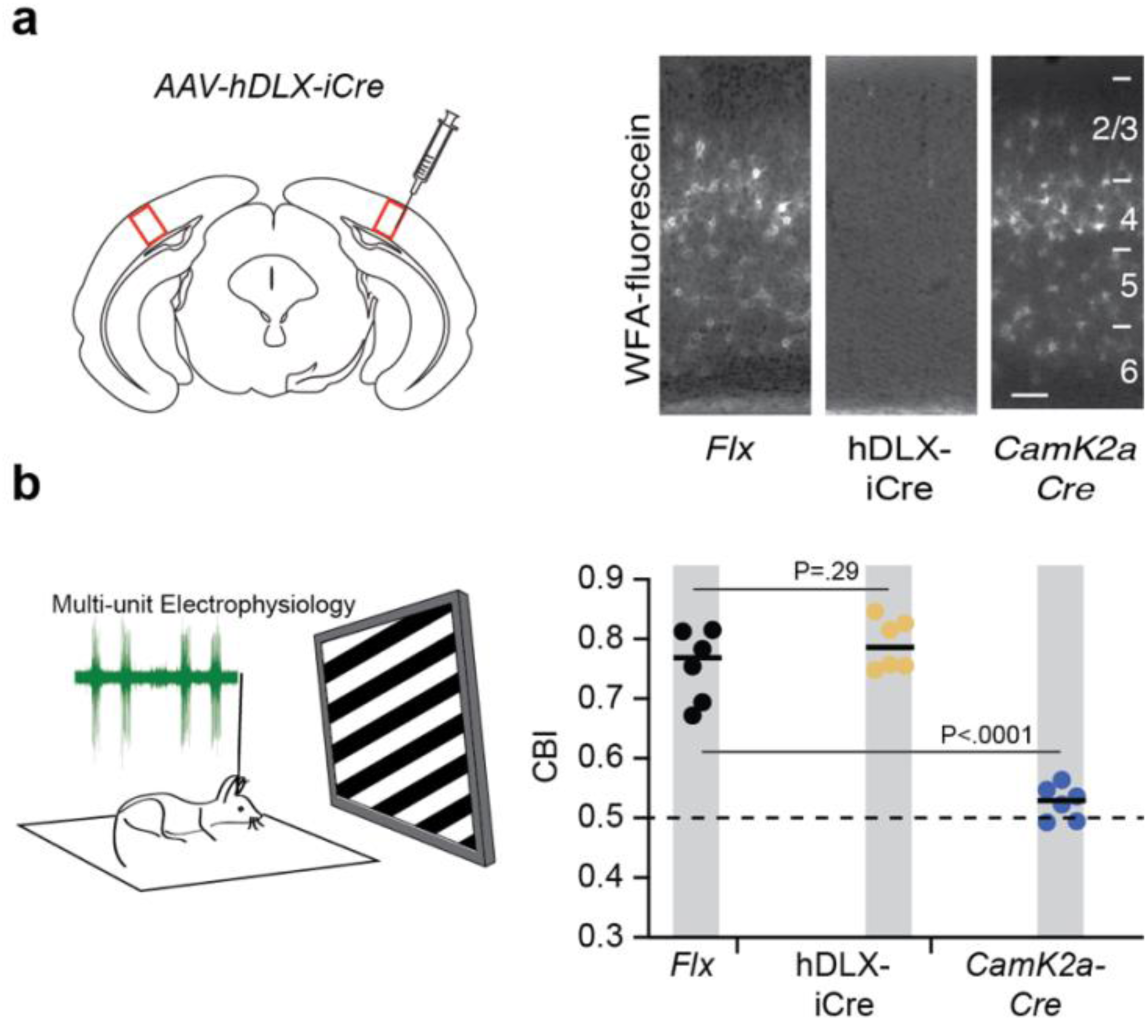
Eliminating *Acan* expression in excitatory cortical neurons does not affect PNNs but prevents the critical period from closing. **(a)** (*left*) Cartoon of a coronal section of mouse brain and injection sites for AAV-hDlx-iCre to delete the *Acan* gene in inhibitory neurons. (*right*) Coronal sections of V1 stained with WFA-fluorescein to label PNNs in both the adjoining hemisphere (*Flx*) and injected hemisphere (hDLX-iCre) of the same mouse and a section from an *Acan flx/flx*; *CamK2a-Cre* mouse. Positions of cortical layers are indicated at right. Scale bar = 100 microns. **(b)** (*left*) Cartoon of the multi-unit recording method for anesthetized mice. (*right*) Mean CBI scores after 4-5d of MD for adult control *Acan flx/flx* mice (*Flx*; n = 6), adult *Acan flx/flx* mice injected with AAV-hDLX-iCre (*hDLX-iCre*, n=6) and adult *Acan flx/flx*;*CamK2a-Cre* mice (*CamK2a-Cre*, n=6). 1-ANOVA.

Given these collective findings, we tested whether deleting expression of *Acan* only in excitatory neurons was sufficient to prevent closure of the critical period. We combined the Acan *flx* allele with *Camk2a-Cre* to delete *Acan* in excitatory neurons in forebrain including V1 (*42*–*44*). Adult *Acan flx/flx; Camk2a-Cre* mice displayed OD plasticity in response to 4-5d MD similar to adult *Acan flx/flx*; *BAF53b-Cre* mice (Fig. 3b). PNNs appeared unaffected by this genetic manipulation (Fig. 3a). Thus, deletion of *Acan* in excitatory neurons was sufficient to prevent the critical period from closing.

### Loss of neuronal Acan elevates expression of some immediate early genes

Last, we assessed how the deletion of *Acan* from inhibitory neurons or all neurons affected the transcriptional profile of PV+ interneurons and surrounding pyramidal neurons that we investigated with electrophysiology and calcium imaging. We performed spatial RNA quantification with the Xenium platform and 5K mouse gene panel on fixed frozen mouse brain sections (*45*). We analyzed 8-9 individual sections from 2 adult male mice (P90-100) for each of the following genotypes: *Acan flx/*flx, *Acan flx/flx; GAD2-Cre*, *Acan flx/flx; CamK2a-Cre*, and *Acan flx/flx; BAF53b-Cre* (Fig 4a). For each section we obtained 15,329 cells on average (ranges 11,530 – 18,305), with 1461 transcript counts and 768 genes per cell on average. We manually circumscribed V1 for each section and annotated the cells using the V1 cortex subset of Allen Brain Atlas single-cell RNA sequencing data as the reference and compared the gene expression profiles for each Cre driver relative to *Acan flx/flx* mice (Fig. 4b)(*46*).

**Figure 4.**
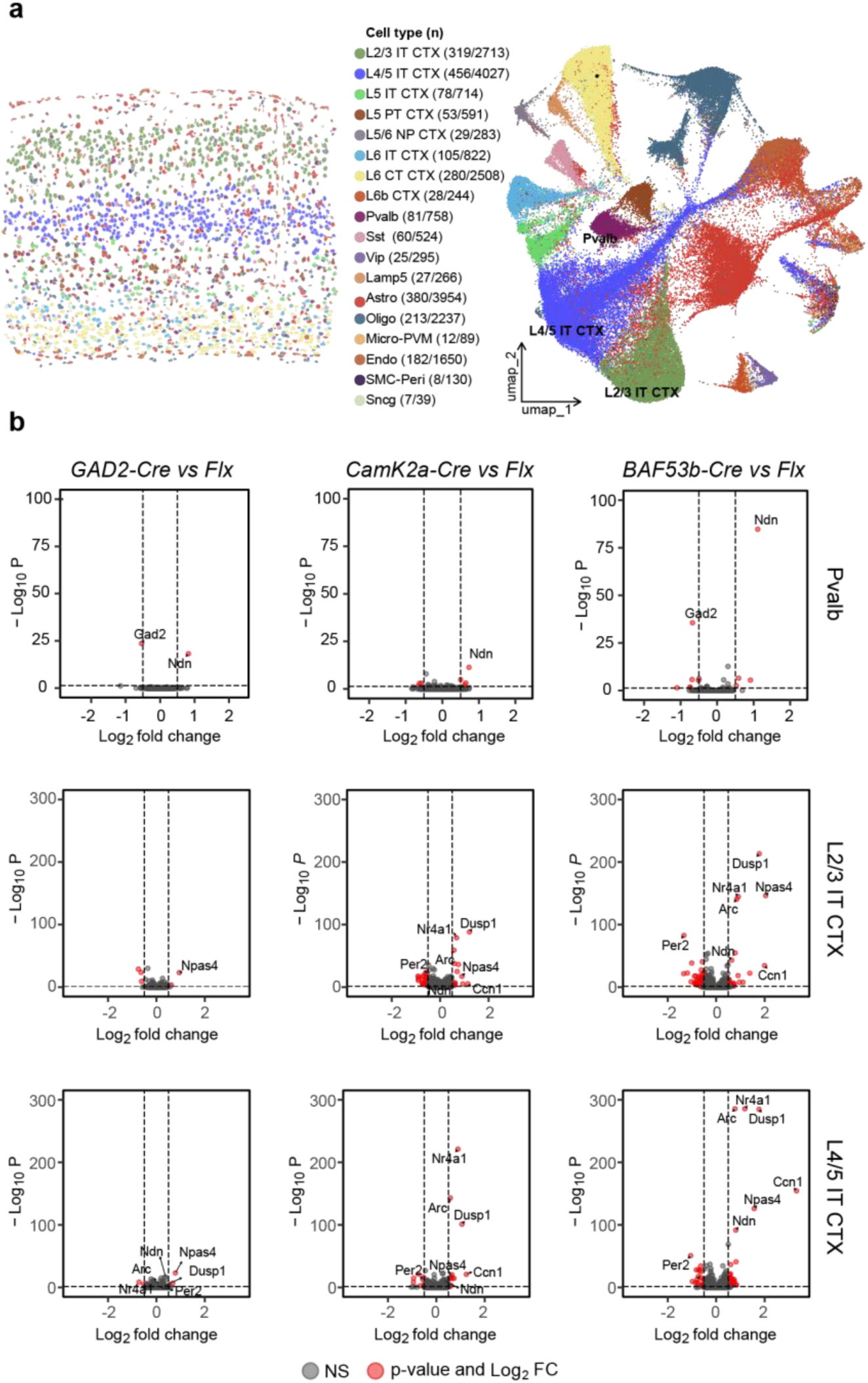
Spatial transcriptomics reveal loss of neuronal *Acan* elevates expression of some immediate early genes. **(a)** (*left*) Segmentation of a representative coronal section of V1 from an adult *Acan flx/flx* mouse and unsupervised clustering followed by cell type label transfer with the Allen V1 Cortex reference dataset. (*middle*) The cell types identified in the data set. Numbers in parentheses indicate the number of cells in V1 of the coronal section / number of cells in V1 in aggregate for *Acan flx/flx* mice. (*right*) The distribution of cell types in UMAP space for sections from *Acan flx/flx* mice. **(b)** Volcano plots for differentially expressed genes when comparing *GAD2-Cre*, *CamK2a-Cre*, and *BAF53b-Cre* versus *Flx* alone for PV+ interneurons (Pvalb), L2/3 IT neurons, and L4/5 IT neurons. Discussed genes with high significance are labeled. NS, non-significant.

There were few differentially expressed genes in V1 between genotypes. Deletion of *Acan* with *GAD2-Cre* resulted in a minor increase in mRNA levels in PV+ interneurons for *Necdin* (*Ndn*) (1.8X), a pleiotropic protein that supports neuronal function and viability (*47*), and a decrease for *Gad2* (0.7X), one of the enzymes that synthesizes the inhibitory neurotransmitter GABA (*48*). However, there was little effect on excitatory neurons in L2/3 or L4/5 save for an increase in expression of the immediate early gene (IEG) *Npas4* (1.5-4X) (Fig. 4b)(*49*). By comparison, deletion of *Acan* with *CamK2a-Cre* yielded a similar increase of *Ndn* in PV+ interneurons (1.6X) and of *Npas4* in L2/3 and L4 excitatory neurons (1.5-1.9X), but this was accompanied by increased mRNA expression of several additional IEGs: *Arc* (1.5X), *Dusp1* (2.0-3.4X), *Nr4a1* (1.6-1.8X), and *Ccn1* (2.3X) (Fig. 4b)(*50*–*53*). Interestingly, the transcriptional profile for PV+ interneurons, L2/3 excitatory neurons, and L4/5 excitatory neurons in *Acan flx/flx; BAF53b-Cre* mice was largely the combination of differentially expressed genes for *Acan* deletion with *GAD2-Cre* and *CamK2a-Cre* (Fig. 4b).

## Discussion

The maturation of PNNs overlaps with the closure of the critical period, but with important caveats. In cats, PNNs are sparse during the zenith of the critical period, increase as the critical period closes, and eventually plateau several months later (*54*, *55*). Dark rearing delays the closure of the critical period but does not prevent the emergence of PNNs; rather dark rearing halves the number of PNNs only in the extragranular layers of V1 (L2/3 and L5) but not in the thalamorecipient granular layer (L4) where PNNs are most numerous (*55*–*57*). By comparison, there are more PNNs in rat V1 during the critical period, amounting to approximately half the total present in adults. Dark rearing again only moderately reduces the number of PNNs in L2/3 and L5 without affecting PNNs in L4 (*15*). Similar to rats, mice display only a small increase (∼20%) in the number of PNNs from the zenith of the critical period to adulthood and the effects of dark rearing are modest (*16*). Despite these similarities between species in the distribution of PNNs across age and visual experience, PNNs are commonly stated to appear with closure of the critical period (*3*, *4*).

Several disparate mechanisms have been proposed for how PNNs may regulate the maturation of brain circuitry. PNNs are proposed serve as extracellular reservoirs for physiologically relevant cations (*58*). They are proposed act as physical barrier that limits the lateral mobility of neurotransmitter receptors as well as synapse formation or loss by PV+ interneurons (*59*, *60*). PNNs also bind a number of growth factors, cell-adhesion molecules, and other factors present in the extracellular environment, and are proposed to regulate structural and/or synaptic plasticity (*7*). To our knowledge, none of these putative mechanisms have been demonstrated to contribute to the closure of the critical period and/or limit visual plasticity.

The strongest functional evidence that implicates PNNs in the closure of the critical period is the phenotype of mice lacking the expression of *Hapln1* in the central nervous system. PNNs are reduced in these mutant mice although the overall levels of many CSPGs are normal and adult mutants retain visual plasticity otherwise confined to the critical period (*5*). However, this study employed visually-evoked potentials (VEPs) to measure OD plasticity, similar to experiments that reported PNNs limit visual plasticity by binding Semaphorin 3A, a chemorepulsive axon guidance protein (*61*). VEPs have limited utility for measuring OD plasticity because this local field potential is not an accurate measurement of neuronal firing activity in V1 (*62*, *63*).

In comparison, we measured OD plasticity with both multi-unit electrophysiologic recordings in anesthetized mice and calcium imaging in awake mice. Multi-unit recordings are the gold standard for measuring OD plasticity (*41*). This technique has been used effectively since the original studies in kittens (*64*). Spiking events detected with high-impedance tungsten electrodes (5-10 mega-ohm) are the basis for the calculation of CBI scores in these experiments (*23*, *24*, *43*, *44*). *In vivo* calcium imaging complements these electrophysiologic recordings by providing cellular resolution of neuronal responses to the visual stimuli presented independently to each eye (*65*). We demonstrate that OD plasticity re-activated by expression of chABC in adult WT mice mirrors OD plasticity in adult mice lacking *Acan* expression in excitatory cortical neurons. Deletion of *Acan* in PV+ interneurons and the loss of PNNs were dispensable for closure of the critical period when tested with 3 different Cre drivers. We conclude that aggrecan outside of PNNs in the surrounding neuropil closes the critical period for OD plasticity.

We probed whether deleting *Acan* either in inhibitory neurons to eliminate PNNs, or in all neurons to remove both PNNs and aggrecan in the neuropil, altered intracortical circuitry. Almost all PV+ interneurons are encapsulated by a discernable PNN (*23*), so we mapped the distribution and strength of excitatory cortical synapses onto PV+ interneurons in L2/3 of V1 with LSPS in adult *Acan flx/flx* mice alone or in combination with *GAD2-Cre* or *BAF53b-Cre*. Most excitatory synapses were from neurons in L4 and did not differ in distribution or strength between *Acan flx/flx mice* and *Acan flx/flx; GAD2-Cre* mice. However, deletion of *Acan* in all neurons with *BAF53b-Cre* resulted in a detectable decrease of the strength of these excitatory synapses. A reduction in intracortical excitatory synaptic drive onto L2/3 PV+ interneurons is also present in mutant mice lacking a functional *nogo-66 receptor 1* gene (*Rtn4r/Ngr1*); *Ngr1* is one of the few other identified genes required to close the critical period for visual plasticity (*24*, *66*, *67*). Interestingly, NGR1 is a synaptic protein and a putative receptor for aggrecan (*68*). This decreased excitatory drive onto PV+ interneurons may tilt balance of excitatory/inhibitory neurotransmission (E/I balance) and faciliate experience-dependent plasticity (*69*).

To assess if the loss of PNNs in *Acan flx/flx; GAD2-Cre* mice or if the reduction in excitatory drive onto PV+ interneurons in *Acan flx/flx; BAF53b-Cre* mice altered the function of excitatory neurons in V1, we measured the response properties of L2/3 excitatory neurons in non-deprived adult mice with calcium imaging. The SNR and average delta F/F were similar between genotypes. PV+ interneurons contribute to the orientation tuning of pyramidal neurons (*70*). Orientation tuning was not perturbed by the deletion of *Acan* with *GAD2-Cre* or *BAF53b-Cre*. These characteristics also resembled those of adult *Ngr1* mutant mice (*34*). However, while the tuning properties of L2/3 excitatory neurons of *Ngr1* mutants are normal, the stability of tuning properties does not increase in adulthood as observed in WT mice (*34*). We suspect this sustained instability of neuronal tuning is the basis for critical-period visual plasticity in adult *Ngr1* mutant mice. Exploring whether adult *Acan flx/flx; BAF53b-Cre* also maintain greater tuning instability is a logical next step.

Interestingly, we detected few differences in the transcriptional profile of cells in adult V1 between genotypes. Genetic deletion of *Acan* in PV+ interneurons has been speculated to yield some manner of compensation (*39*). We find no convincing evidence in our spatial transcriptomic analysis to support this conjecture. The mRNA expression of only two genes altered in V1 of *Acan flx/flx; GAD2-Cre* mice, *Ndn* and *Gad2,* and these differences were modest in magnitude. Moreover, these genes exhibited similar transcriptional changes in *Acan flx/flx; BAF53b-Cre* mice that retained critical-period OD plasticity in response to brief MD while *Acan flx/flx; GAD2-Cre* mice did not. By comparison, *Acan flx/flx; CamK2a-Cre* and *Acan flx/flx; BAF53b-Cre* both displayed increased mRNA expression of several IEGs.

We suspect the elevated expression of these IEGs by excitatory neurons in L2/3 and L4 are a consequence of sustained developmental plasticity. Overexpression of *Arc* has been reported to sustain critical-period OD plasticity in adult mice, but this involved ∼5X increase in mRNA expression and was measured with VEPs (*71*). By comparison, we detected a ∼1.75X increase in *Arc*. Both the *Dusp1* gene encoding Dual specificity protein phosphatase 1, and *Nr4a1* gene, encoding an ‘orphan’ nuclear hormone receptor, are expressed widely throughout the body and are elevated by stress responses across numerous cell types (*51*, *53*). Neither appears to have been previously implicated in visual plasticity. *Ccn1* encodes cellular communication network factor 1, a secreted protein that was first identified as a growth factor IEG in fibroblasts and is critical for vascular development (*52*, *72*). A recent study reported that *Ccn1* stabilizes the tuning properties of excitatory neurons in mouse V1 and attenuates OD plasticity following 4d of MD in adult mice (*73*). In contrast, we and others do not observe OD plasticity in adult WT mice following 4d of MD (*21*, *34*, *74*), and here we detect a 4-10X increase in *Ccn1* mRNA in *BAF53b-Cre* mice that display sustained critical-period plasticity.

In summary, evidence for PNNs closing the critical period is predominantly circumstantial and often mischaracterized. Many consequences of treatment with chABC have been misattributed to PNNs although this enzyme not only degrades PNNs, but also CSPGs and other components of the extracellular matrix in the surrounding neuropil. Here we determined that PNNs are dispensable for closing the critical period. We propose that chABC re-activates visual plasticity otherwise confined to the critical period by degrading aggrecan expressed by excitatory cortical neurons. This sustained developmental plasticity is associated with a reduction in excitatory synaptic drive onto PV+ interneurons but lacks an obvious transcriptional profile that pinpoints how aggrecan may limit visual plasticity. Future work will be required to discover the mechanism by which aggrecan operates to close the critical period, as well as to assess how these findings inform other experimental paradigms proposed to be regulated by PNNs such as fear conditioning and spatial memory (*3*, *4*).

## Methods

| REAGENT or RESOURCE | SOURCE | IDENTIFIER |
| --- | --- | --- |
| **Deposited data** |  |  |
| Calculated ocular dominance for all units from multi-unit electrophysiologic recordings |  | N/A |
| Calculated tuning properties for mice and neurons from calcium imaging |  | N/A |
| Calculated synaptic responses for all neurons for LSPS |  | N/A |
| Behavioral data for visual acuity measurements |  | N/A |
| Xenium transcriptomic data |  | N/A |
| **Experimental models: Organisms/strains** |  |  |
| Acan flx | Rowlands et al. 2018 |  |
| PV-Cre | The Jackson Laboratory | RRID: ISMR_JAX: 017320 |
| Dlx5/6-Cre | The Jackson Laboratory | RRID: ISMR_JAX: 008199 |
| GAD2-Cre | The Jackson Laboratory | RRID: ISMR_JAX: 028867 |
| BAF53b-Cre | The Jackson Laboratory | RRID: ISMR_JAX: 027826 |
| CamK2a-Cre | The Jackson Laboratory | RRID: ISMR_JAX: 005359 |
| Mouse: B6;DBA-Tg(tedO-GCaMP6s)2Niell/j | The Jackson Laboratory | RRID: ISMR_JAX: 024742 |
| Mouse: B6;Cg-Tg(Camk2a-tTA)1Mmay/DboJ | The Jackson Laboratory | RRID: ISMR_JAX: 007004 |
| **Software and algorithms** |  |  |
| MATLAB | Mathworks | https://www.mathworks.com/ |
| Processing2 | Processing | https://processing.org/ |
| ImageJ | NIH | RRID:SCR_003070 |
| Xenium Analyzer Onboard Analysis | 10X Genomics 4.0.1.4 | RRID:SCR_026158 |
| Xenium Explorer | 10X Genomics 4.1.1 | RRID: SCR_025847 |
| **Reagents** |  |  |
| WFA-fluorescein | Thermofisher | RRID:AB_2315605 |
| Anti-Parvalbumin antibody | R&D systems | RRID:AB_2173907 |
| R-CPP | Tocris | RRID:SCR_003689 |
| MNI-glutamate | Tocris | RRID:SCR_017300 |
| Normal horse serum | Vector Labs | RRID:AB_2336617 |
| DAPI | Thermofisher | RRID:AB_2629482 |
| AAV-mDLX-GFP | Addgene | RRID:Addgene_83900 |
| AAV-hDLX-Cre | Addgene | RRID:Addgene_164450 |

### Lead contact

Further information and requests for resources and reagents should be directed to and will be fulfilled by the Lead Contact, Aaron McGee

### Data availability

The data listed in the table above have been deposited in Mendeley data.

### Experimental model and subjects

All procedures were approved by University of Louisville and University of Arizona Institutional Animal Care and Use Committees and were in accordance with guidelines set by the US National Institutes of Health. Mice were anesthetized by isoflurane inhalation and euthanized by carbon dioxide asphyxiation or cervical dislocation following deep anesthesia in accordance with approved protocols. Mice were housed in groups of 5 or fewer per cage in a 12/12 light dark cycle. Animals were naive subjects with no prior history of participation in research studies.

### Mice

The *Acan flx* allele was a generous gift from Gunnar Dick at the University of Oslo, Norway(*19*). The Cre Drivers *PV-Cre* (strain# 008069), *Dlx5/6-Cre* (strain# 008199), *GAD2-Cre* (strain# 019022), *BAF53b-Cre* (strain# 027826), and *CamK2a-Cre* (strain# 005359), were obtained from Jackson Labs(*23*, *44*). Calcium imaging was performed on mice expressing GCaMP6S in excitatory neurons in forebrain. The *CaMKII-tTA* (strain# 007004) and *TRE-GCaMP6s* (strain# 024742) transgenic mouse lines were also obtained from Jackson Labs(*75*, *76*). Mice were genotyped with primer sets suggested by Rowlands et al. (2018) and Jackson Labs(*19*).

### Histology of visual cortex

Sectioning and immunostaining were performed as described previously(*23*). In brief, mice were deeply anesthetized with Ketamine HCl (200mg/Kg, Zoetis)/Xylazine (20mg/Kg, Covetrus) and transcardially perfused with phosphate buffered saline (PBS; Alkali Scientific) followed by a PBS buffered 4% paraformaldehyde (PFA) (Thermo Scientific). Brains post-fixed overnight in 4% PFA/PBS. Free-floating 60-micron sections were cut on a vibrating microtome (Leica VT 1000S) and preserved in PBS containing 0.05% sodium azide (Fisher).

Coronal sections containing visual cortex were incubated in blocking solution, 3% normal horse serum (NHS; Vector Laboratories S-2000) in Tris-buffered saline (TBS) containing 0.1% Triton X-100 (Sigma-Aldrich) for 1 hour at room temperature (TBS-T). In sections labeled for perineuronal nets (PNNs), fluorescein conjugated Wisteria Floribunda Agglutinin (WFA) (VectorLabs, FL-1351) was diluted in blocking solution to at 2mg/mL. In sections double labeled for PNNs and parvalbumin (PV), both WFA and the primary antibody sheep anti-PV (R&D Systems, AF5058) were diluted in blocking solution to 1mg/mL. Sections were incubated at room temperature (RT) for 1 hour. After repeated washing in TBS-T (3 X 10 min), sections for double labeling were incubated in secondary antibody, 594-conjugated donkey anti-sheep (Jackson Immuno Research) 1:200 in blocking solution, for 1 hour at room temperature. After a final series of washes (3x 10 min in TBS-T, 1 X 10 min in TBS), sections were washed with TBS containing 4’,6-diamidino-2-phenylindole (DAPI) at 1μg/ml (Invitrogen) for 1 X 10 min, followed by another final wash in TBS alone. Sections were mounted onto SuperFrost Plus slides (Fisher).

### Analysis of PNN number

Images from coronal sections stained with WFA-fluorescein were captured with a BX-51 microscope, 20x 0.4 NA objective and 12-bit monochrome camera (Retiga EX, QImaging). DAPI staining was utilized to identify visual cortex prior to capturing images of PV/WFA density. Two images were required to span the distance from the subcortical white matter to the pial surface. Images were merged with the software Photoshop following linear contrast adjustment. Data points are the average of at least three sections from each of three animals for each genotype.

### Monocular deprivation

One eye lid was sutured shut between postnatal day 26-28 with 6-0 polypropylene monofilament (Prolene 8709H; Ethicon) under brief isoflurane anesthesia for 4-5 days. The knot was sealed with cyanoacrylate glue. Upon removing the suture, the eye was examined under a stereomicroscope and mice with scarring of the cornea were eliminated from the study.

### Multi-unit electrophysiologic recordings

Recordings and analysis were performed blind to treatment. Methods were adapted from previously published methods(*43*, *44*). In brief, mice were anesthetized with isoflurane (4% induction, 1-2% maintenance in O2 during surgery). The mouse was placed in a stereotaxic frame, and body temperature was maintained at 37°C by a homeostatically-regulated heat pad (TCAT-2LV, Physitemp). Dexamethasone (4 mg/kg s.c.; American Reagent) was administered to reduce cerebral edema. The eyes were flushed with saline, and the corneas were protected thereafter by covering the eyes throughout the surgical procedure with silicone oil (10838, Millipore Sigma). A craniotomy was made over visual cortex in the left hemisphere, and a custom-designed aluminum head bar was attached with Metabond over the right hemisphere to immobilize the animal during recording. Prior to transfer to the recording setup, a dose of chlorprothixene (0.5 mg/kg i.p.; C1761, Sigma) was administered to decrease the level of isoflurane required to maintain anesthesia to 0.8%.

Recordings were made with Epoxylite-coated tungsten microelectrodes with tip resistances of 5-10 mega-ohm (FHC). The signal was amplified (model 3600; A-M Systems), low-pass filtered at 3000Hz, high-pass filtered at 300Hz, and digitized (micro1401; Cambridge Electronic Design). Multi-unit activity was recorded from 3-6 locations separated by >90μm in depth for each electrode penetration. In each mouse, each penetration separated by at least 200 microns and were distributed across the binocular region of primary visual cortex (V1), defined by a receptive field azimuth <25°. Responses were driven by drifting sinusoidal gratings (0.1cpd, 95% contrast), presented in six orientations separated by 30° (custom software, MATLAB). The gratings were presented for 1s of each 3s trial. The grating was presented in each orientation in a pseudorandom order at least four times, interleaved randomly by a blank, which preceded each orientation once. Action potentials (APs) were identified in recorded traces with Spike2 (Cambridge Electronic Design). Only waveforms extending beyond 4 standard deviations above the average noise were included in subsequent analysis. For each unit, the number of APs in response to the grating stimuli was summed and averaged over the number of presentations. If the average number of APs for the grating stimuli was not greater than 50% above the blank, the unit was discarded.

The Ocular Dominance Index (ODI) was calculated as (C-I)/(C+I), where C (contralateral) and I (ipsilateral) are the average number of APs elicited in a given unit when presenting the same visual stimulus to each eye independently. Units were then assigned to one of seven OD categories (1-7) where units assigned to category 1 are largely dominated by input from the contralateral eye, and units assigned to category 7 are largely dominated by input from the ipsilateral eye.

### Calcium imaging to measure neuronal tuning properties

Imaging and analysis were performed blind to genotype. Methods were identical to our previously published experiments(*35*). Wide-field (epifluorescent) and cellular resolution (two-photon laser scanning) imaging experiments were performed though a cranial window. In brief, mice were administered carprofen (5 mg/kg, Zoetis) and buprenorphrine-XR (0.1 mg/kg, Fidelis) for analgesia and anesthetized with isoflurane (4% induction, 1-2% maintenance). The hair on the scalp was clipped and mice were mounted on a stereotaxic frame with palate bar and their body temperature maintained at 37°C with a heat pad controlled by feedback from a rectal thermometer (TCAT-2LV, Physitemp). The scalp was resected, the connective tissue removed from the skull, and an aluminum headbar affixed with C&B metabond (Parkell). A circular region of bone 3mm in diameter centered over left visual cortex was removed using a high-speed drill (Foredom). Care was taken to not perturb the dura. A sterile 3mm circular glass coverslip was sealed to the surrounding skull with cyanoacrylate (Pacer technology) and dental acrylic (Ortho-jet, Lang Dental). The remaining exposed skull likewise sealed with cyanoacrylate and dental acrylic. Mice recovered on a heating pad. Mice were left to recover for at least 2 days prior to imaging.

After implantation of the cranial window, the binocular zone of visual cortex was identified with wide-field calcium imaging similar to our method for optical imaging of intrinsic signals(*22*). In brief, mice were anesthetized with isoflurane (4% induction), provided a low dose of the sedative chlorprothixene (0.5mg/kg i.p.; Sigma) and secured by the aluminum headbar. The eyes were lubricated with a thin layer of ophthalmic ointment. Body temperature was maintained at 37°C with heating pad regulated by a rectal thermometer (TCAT-2LV, Physitemp). Visual stimulus was provided through custom-written software (MATLAB, Mathworks). A monitor was placed 25cm directly in front of the animal and subtended +40 to -40 degrees of visual space in the vertical axis. A horizonal white bar (2 degrees high and 20 degrees wide) centered on the zero-degree azimuth drifted from the top to bottom of the monitor with a period of 8s. The stimulus was repeated 60 times. Cortex was illuminated with blue light (475 ± 30nm) (475/35, Semrock) from a stable light source (Intralux dc-1100, Volpi). Fluorescence was captured utilizing a green filter (HQ620/20) attached to a tandem lens (50mm lens, Computar) and camera (Manta G-1236B, Allied Vision). The imaging plane was defocused to approximately 200 microns below the pia. Images were captured at 10Hz as images of 1024x1024 pixels and 12-bit depth. Images were binned spatially 4x4 before the magnitude of the response at the stimulus frequency (.125 Hz) was measured by Fourier analysis.

For imaging at cellular resolution, mice were mounted by the headplate atop a spherical treadmill. The monitor was centered on the zero azimuth and elevation 35cm away from the mouse and subtended 45 (elevation) by 80 degrees (azimuth) of visual space. A battery of static sinusoidal gratings were generated in real time with custom software (Processing, MATLAB) as described(*77*). Stimulus presentation was synchronized to the imaging data by time stamping the presentation of each visual stimulus to the image acquisition frame number a transistor-transistor logic (TTL) pulse generated with an Arduino at each stimulus transition. Orientation was sampled at equal intervals of 30 degrees from 0 to 150 degrees (6 orientations). Spatial frequency (SF) was sampled in 8 steps on a logarithmic scale at half-octaves from 0.028 to 0.48 cycles per degree (cpd). An isoluminant grey screen was included (blank) as a 9th step in the SF sampling as a control. Spatial phase was equally sampled at 45-degree intervals from 0 to 315 degrees for each combination of orientation and SF. Gratings with random combinations of orientation, SF, and spatial phase were presented at a rate of 4Hz on a monitor with a refresh rate of 60Hz. Imaging sessions were 10 minutes (2400 presentations in total). Consequently, each combination of orientation and SF was presented 40 times on average (range 29-56).

Imaging was performed with a resonant scanning two-photon microscope controlled by Scanbox image acquisition and analysis software (Neurolabware). The objective lens was fixed at vertical for all experiments. Fluorescence excitation was provided by a tunable wavelength infrared laser (Ultra II, Coherent) at 920 nm. Images were collected through a 16x water-immersion objected (Nikon, 0.8 NA). Images (512x796 pixels, 520x740 microns) were captured at 15.5 Hz at depths between 150 – 400 microns. Eye movements and changes in pupil size were recorded using a Dalsa Genie M1280 camera (Teledyne Dalsa) fitted with 50 mm 1.8 lens (Computar) and an 800nm long-pass filter (Edmunds Optics). Imaging was performed on alert mice positioned on a spherical treadmill by the aluminum head bar affixed to the skull. The visual stimulus was presented to each eye separately by covering the fellow eye with a small custom occluder.

The imaging series for each eye were motion corrected with the SbxAlign tool. Regions of interest (ROIs) corresponding to excitatory neurons were selected manually with the SbxSegment tool following computation of pixel-wise correlation of fluorescence changes over time from 350 evenly spaced frames (∼4%). ROIs for each experiment were determined by correlated pixels the size similar to that of a neuronal soma. The fluorescence signals for each ROI and the surrounding neuropil were extracted from this segmentation map.

The fluorescence signal for each neuron was extracted by computing the mean of the calcium fluorescence within each ROI and subtracting the median fluorescence from the surrounding perimeter of neuropil(*77*, *78*). An inferred spike rate (ISR) was estimated from adjusted fluorescence signal with the Vanilla algorithm(*79*). A reverse correlation of the ISR to stimulus onset was used to calculate the preferred stimuli (*77*, *78*, *80*, *81*). Neurons that satisfied the following three criteria were categorized as visually responsive: (1) the ISR was highest with the optimal delay of 4-9 frames following stimulus onset. This delay was determined empirically for this transgenic GCaMP6s mouse(*77*); (2) the SNR was at least one standard deviation greater than spontaneously active neurons. The signal is the mean of the spiking standard deviation at the optical delay between frames 4-9 after stimulus onset and the noise this value at frames -2 to 0 before the stimulus onset or frames 15-18 after it (*77*, *80*); (3) and the percent of responses to the preferred stimulus was at least one standard deviation greater than spontaneously active neurons. Visual responsiveness for every neuron was determined independently for each eye. The visual stimulus capturing the preferred orientation and SF was the determined from the matrix of all orientations and SFs presented as the combination with highest average ISR.

The preferred orientation for each neuron was calculated as:

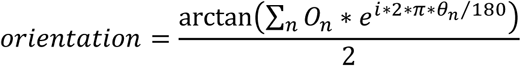

O_n_ is a 1x6 array of the mean z-scores associated with the calculation of the ISR at orientations O_n_ (0 to 150 degrees, spaced every 30 degrees). Orientation calculated with this formula is in radians and was converted to degrees.

The preferred SF for each neuron was calculated as:

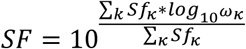

Sfk is a 1x8 array of the mean z-scores at SFs wk (8 equal steps on a logarithmic scale from 0.028 to 0.481 cpd). Tails of the distribution were clipped at 25% of the peak response. The tuning width was the full width at half-maximum of the preferred SF in octaves. The percent of visually responsive neurons with significant responses at each SF was determined by comparing the distribution of ISR values at each SF versus the stimulus blank with a KW-test with Dunn’s correction for 8 comparisons. Neurons with P<.01 for a given SF were considered significant responses at that SF(*82*).

Orientation half-maximum full widths were calculated by linear interpolation to determine the position (in degrees) corresponding to half of the maximum responses of the ISR. Binocular matching was measured as the absolute difference in the preferred orientation calculated for visual stimuli presented to the contralateral eye and ipsilateral eye along the 180 degree cycle(*83*–*85*). Neuronal ODI was calculated as (C-I)/(C+I), where C and I are the mean normalized change in fluorescence (dF/F) for the preferred visual stimulus for the contralateral eye (C) and ipsilateral eye (I), respectively. In cases where neurons displayed no significant response to visual stimuli provided to one eye, they were considered monocular for the other eye and assigned ODI values of 1 (contralateral) and -1 (ipsilateral)(*82*). Summed ODI was calculated by summing the dF/F for the preferred visual stimulus for the neurons visually responsive to the contralateral eye (C) and ipsilateral eye (I) for each mouse, respectively. The summed ODI per mouse was then calculated as (C-I)/(C+I) for each mouse.

### Patch clamp electrophysiology and laser scanning photostimulation (LSPS)

Electrophysiology recording and LSPS mapping experiments were performed on postnatal day ∼60. Mice were deeply anesthetized by isoflurane and quickly decapitated. The visual cortex was sectioned into coronal slices of 350 mm thickness using a vibratome (VT1200S; Leica Systems) in ice-cold artificial cerebrospinal fluid (ACSF) (in mM: 126 NaCl, 2.5 KCl, 26 NaHCO3, 2 CaCl2, 2 MgCl2, 1.25 NaH2PO4, and 10 glucose). The slices were then incubated in ACSF saturated with 95% O_2_ and 5% CO_2_ for 30 min at 32 °C and switched to room temperature in the same ACSF for at least 30 min before transferring to the recording chamber.

Slices were visualized with a microscope (BX51WI; Olympus) equipped with infrared differential interference contrast optics and an epifluorescent light source. V1 binocular zone was identified based on the coordinates, as well as laminar and cytoarchitectonic features(*86*), and electrophysiological recordings performed as described previously(*43*). Slices were visualized either with a low magnification objective (4X, NA 0.13, Olympus, for LSPS mapping) or high magnification objective (60X water immersion, LUMPlanFl, NA 0.9 Olympus, for targeted patching). Digital slice images were acquired with a charge coupled digital camera (Retiga 2000DC, Qimaging) and were used for registering photostimulation locations. The GFP-positive PV+ interneurons in binocular zone were identified under DIC and epifluorescence for patch clamp whole-cell recording. Series resistance (Rs) was monitored throughout recordings, only stable (<15% change) cells with Rs < 30 MΩ throughout the recordings were included. Intrinsic properties were measured in current-clamp mode immediately by injecting current steps (1-sec duration, −100 to +500 pA in 50 pA increments). Passive membrane properties were calculated based on current responses to a negative voltage command pulse. Traces were analyzed off-line to identify APs and to calculate the firing frequency–current relationships. Electrophysiological signals were amplified with a Multiclamp 700B amplifier (Molecular Devices) and acquired with BNC 6259 data acquisition boards (National Instruments) and the Ephus software. Signals were digitized and acquired at 10 kHz. The patch electrode was pulled from borosilicate glass and had electrical resistance of 4-6 mega-ohm. For LSPS mapping experiments, the electrode internal solution contains (in mM) 130 K-gluconate, 4 KCl, 2 NaCl, 10 HEPES, 4 ATP-Mg, 0.3 GTP-Na and 14 phosphocreatine (pH 7.2, 295 mOsm).

LSPS mapping on slices were carried out in a perfusion chamber mounted on a motorized stage (Sutter Instruments) at room temperature. The chamber was perfused with modified ACSF with higher concentrations of magnesium and calcium (in mM: 126 NaCl, 2.5 KCl, 26 NaHCO_3_, 4 CaCl_2_, 4 MgCl_2_, 1.25 NaH_2_PO_4_, and 10 glucose) at 1.5-2 ml/min(*87*). This modified ACSF also contains 5 μM R-3-(2-Carboxypiperazin-4-yl)-propyl-1-phosphonic acid (R-CPP) (Tocris/R&D systems) to block NMDA-receptor currents and reduce plasticity and 0.2mM 4-methoxy-7-nitroindolinyl (MNI)-caged glutamate (Tocris/R&D systems). Neuronal cell bodies were at least 50 microns below the surface of the slice. LSPS was performed through a 4× objective lens (NA 0.13, Olympus). 20-mW, 1 millisecond ultraviolet (UV) laser (DPSS Lasers) pulses ((350nm) were scanned onto the sample after passing an electro-optical modulator (Conoptics) and a mechanical shutter (Uniblitz). We have empirically determined that LSPS-evoked APs can only be recorded from stimulation locations within 100 microns of the soma, suggesting that stimulation is rather focal. For LSPS mapping experiments, synaptic currents in patched neurons were detected under voltage clamp. A stimulus grid (16 X 16, 75-micron spacing) was overlaid on the binocular V1 region, spanning from pia to white matter. For each L2/3 PV+ interneuron, the grid was centered horizontally over the soma and aligned at the top edge with the pia. PV+ interneurons were voltage clamped at -70mV (calculated reversal potential for chloride ions) during the LSPS mapping to minimize the contamination of chloride currents.

### Visual water task for measuring acuity

The Visual Water Task was employed to measure acuity(*88*). Training and testing were performed as previously described(*23*, *44*). In brief, two monitors were positioned at the wide end of a trapezoidal tank behind clear plexiglass. One monitor displayed a sinusoidal SF grating at 95% contrast, while the other displayed an isoluminant grey screen. The luminance of the two monitors was matched and gamma corrected with computer software (Eye-One Match 3). Inside the tank, the monitors were separated by a 46cm divider. The SF was determined relative to the length of this divider. The tank was filled with water and a hidden platform submerged below the surface of the water in front of the monitor displaying the grating.

Using a low SF (0.1 cpd), mice were trained to swim towards the monitor displaying the grating and hidden platform after a molding phase during which mice gradually learned to swim from a release chute at the back of the tank towards the monitors. During the training phase, when a mouse chose incorrectly, it repeated the trial on the same side until it chose correctly and was then returned to its home cage. For both the training, and the subsequent testing phase, mice swam blocks of 10 interleaved trials in groups of 5 for a maximum of 4 blocks of trials per day.

During the testing phase, the SF was increased in small, sequential increments until an animal consistently fell to 70% accuracy. Starting at 0.1 cpd, mice had to succeed at three consecutive trials before proceeding to the next special frequency, which presented one more complete cycle of the sinusoidal grating. Following the first failure, mice were required to achieve 5 correct trials in a row, or 8 correct trials out of 10 at each SF before proceeding to the next higher frequency. Once a mouse failed to complete 8 correct trials out of 10 at a given SF, it was briefly retrained at half that SF to eliminate any potential ‘side bias’. Then, testing resumed at the SF below the original failure. The threshold for visual acuity was established once a mouse exhibited a consistent pattern of performance. Acuity thresholds were estimated as the SF average from three or more failures at adjacent SFs. Throughout the testing phase, any mouse that failed to find the hidden platform on the first try repeated the trial one more time before it was returned to its home cage, regardless of whether it chose correctly the second time.

### Xenium spatial transcriptomics and analysis

Mouse visual cortex tissue sections were profiled using the 10x Genomics Xenium *in situ* platform. Briefly, fixed frozen tissue sections were processed according to the Xenium workflow, including probe hybridization, enzymatic amplification, fluorescent imaging, decoding and quantification of targeted transcripts, and cell segmentation. The resulting Xenium Onboard Analysis output included cell-level transcript count matrices, spatial coordinates, and cell segmentation boundaries. ROIs corresponding to V1 sections were defined in Xenium Explorer for downstream analysis, and cells within these regions were retained for data analysis.

Xenium spatial transcriptomics data were analyzed in R using Seurat(*89*). Pipeline-generated Xenium output files were loaded using *LoadXenium* with the Xenium assay, cell segmentations, and spatial coordinates. ROI cell statistics files exported from Xenium Explorer were used to subset cells by section. Initial cell filtering retained cells with transcript count > 10 and number of genes > 5. Additional quality control filtering was performed per section using a 2-time adaptive median absolute deviation-based filter on transcript count and number of genes. For downstream analysis, the Xenium assay was log-normalized with Seurat *NormalizeData*. The top 2,000 variable features were selected using the vst method, and data were scaled across the selected variable features. Principal component analysis was performed, followed by Harmony-based integration across sections using *IntegrateLayers* with the first 20 principal components. A shared nearest-neighbor graph was constructed from the Harmony-integrated reduction using the first 20 dimensions, and cells were clustered with Seurat at resolution 0.6. Uniform Manifold Approximation and Projection (UMAP) embedding was also computed from the first 20 Harmony dimensions.

Cell-type annotation was performed by label transferring from the V1 subset of Allen Brain Atlas mouse single-cell RNA sequencing reference dataset(*46*). The reference object was analyzed using the RNA assay and analyzed to match the Xenium query workflow. Variable features were identified in the reference and intersected with variable genes present in the Xenium data. Transfer anchors were identified using *FindTransferAnchors*, with the first 20 dimensions. Cell subclass labels from the reference metadata were transferred to Xenium cells using *TransferData*. Differential expression analysis between experimental groups was performed using Seurat *FindMarkers* on the annotated cell types. We employed a Model-based Analysis of Single-cell Transcriptomics (MAST) model as the statistical test for the significance of difference(*90*). For each comparison, we retained genes expressed in at least 10% cells of either group. Significant differentially expressed genes (DEGs) were defined using absolute log_2_ fold change > 0.5 and false discovery rate (FDR) adjusted P value < 0.05.

## Acknowledgements

We thank D. Ringach (UCLA) and J. Trachtenberg (UCLA) for sharing software and hardware design for visual stimulus presentation and image analysis prior to publication, A. Eliasen and G. Armstrong for software development, and B. Croslin for mouse husbandry and genotyping. This research is supported by the National Eye Institute (R01EY035138 to SQ and AWM)

## Author Contributions

The contributions are as follows: ECC, TCB, HM, SQ and AWM designed the study. ECC, TCB, XM, HM, KRK, KB, and PC performed the experiments. ECC, TCB, XM, HM, KRK, KB, PC, SQ and AWM analyzed the data. ECC, TCB, HM, PC, SQ and AWM built the figures and wrote the manuscript.

## Declaration of Interests

The authors declare no competing interests

## Notes

### Competing Interest Statement

The authors have declared no competing interest.

### Summary of Updates

Figures in original uploaded version were poor resolution following the conversion to PDF. The original PDF is now provided

