## Supplemental Figures for "Neuropil aggrecan, not perineuronal nets, closes the critical period for visual plasticity"

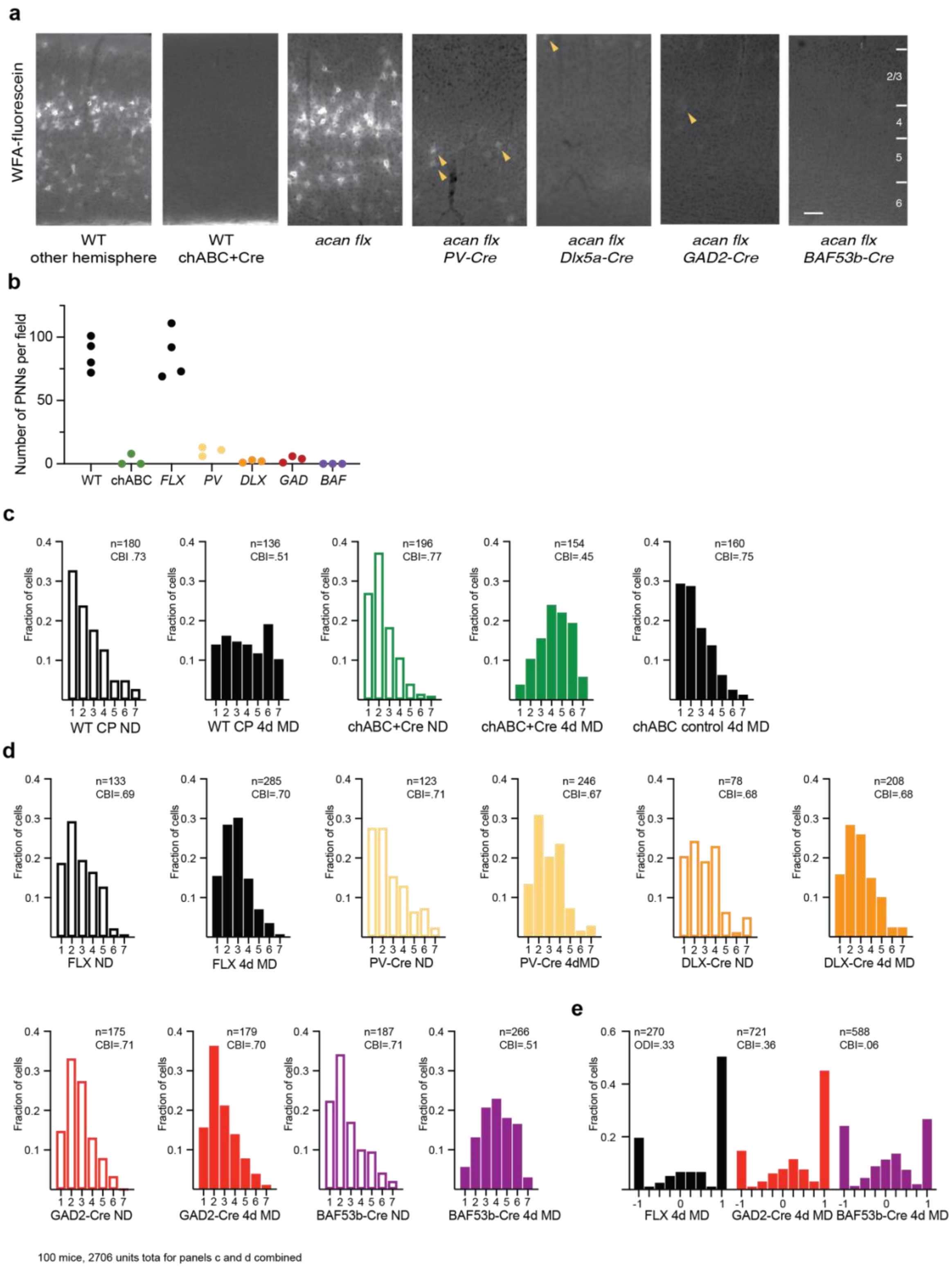


**Figure S1. Deletion of *Acan* gene expression in inhibitory neurons eliminates PNNs but pan-neuronal deletion is required to sustain plasticity similar to cortical expression of chABC. (a)** Larger field of view of coronal sections of visual cortex stained with WFA-fluorescein to label PNNs for both hemispheres of mice injected with AAVs to express chABC (chABC + Cre), and *Acan flx/flx* mice alone and in combination with *PV-Cre*, *Dlx5/6-Cre*, *GAD2-Cre*, or *BAF53b-Cre*. Cortical layers indicated at right. Scale bar = 100 microns **(b)** Quantification of the number of PNNs per imaging field for mice presented in panel a. **(c)** OD histograms for the groups presented in Figure 1 panel b. CBI is the mean of n = the number of units. ND, non-deprived. **(d)** OD histograms for the groups presented in Figure 1 panel d. CBI is the mean of the number of units (n). ND, non-deprived. Panels c and d represent data from 100 mice and 2706 units. **(e)** Histograms of ODI values for neurons *Acan flx/flx* mice alone and in combination with *GAD2-Cre* or *BAF53b-Cre*.

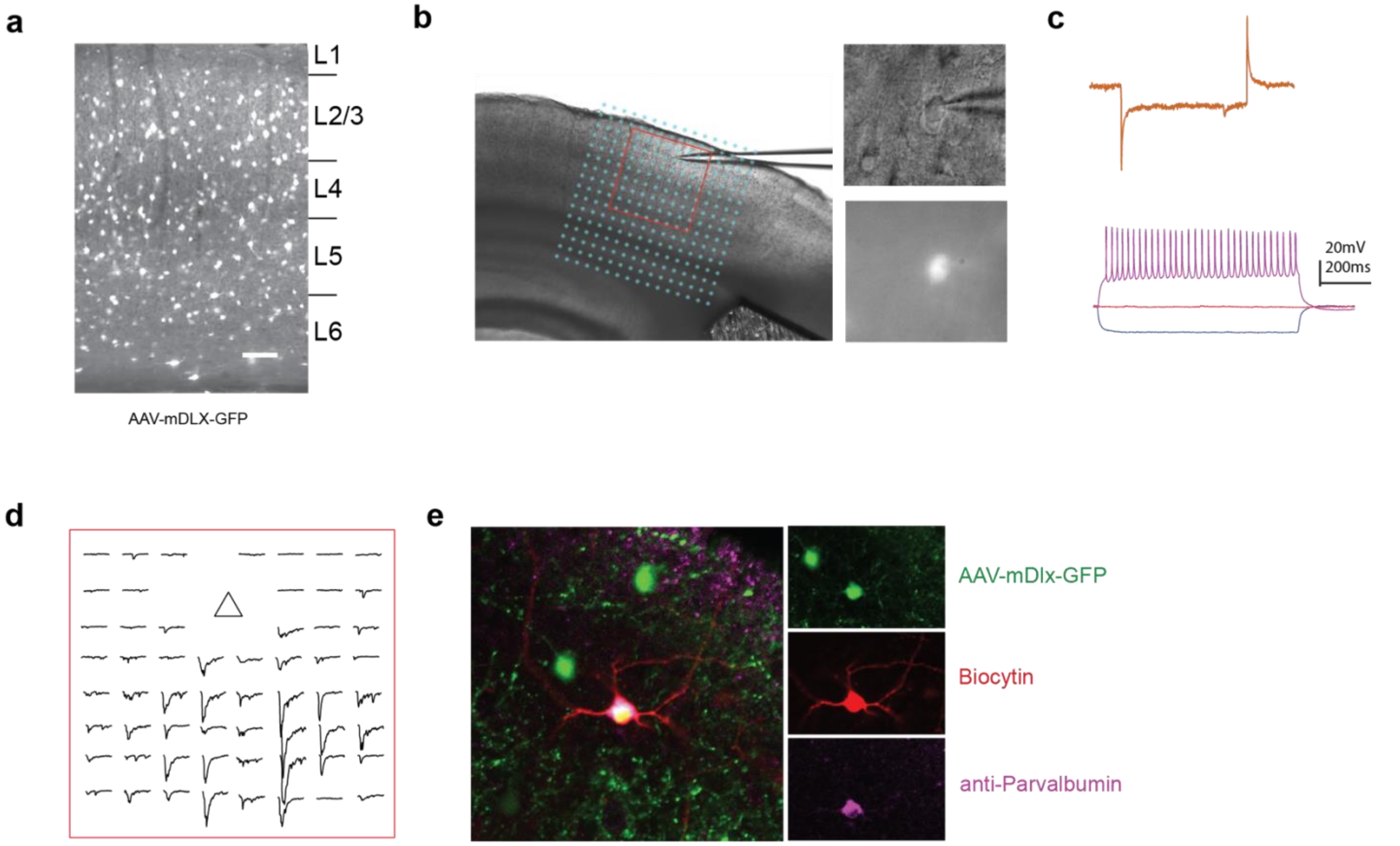


**Figure S2. Validation of mapping excitatory synaptic inputs onto PV interneurons with LSPS. (a)** A coronal section of V1 from an *Acan flx/flx* mouse transduced with AAV-mDLX-GFP-Fishell-1 that labels inhibitory neurons. Cortical layers indicated at right. Scale bar = 100 microns **(b)** (*left*) Example whole-cell recording from a GFP-positive neurons overlaid with the 16X16 grid for glutamate uncaging. (*upper right*) Differential interference contrast image of the neuronal soma and pipet tip. (*lower right*) Fluorescence image of the same cell. **(c)** (*top*) Current response of the L2/3 PV+ interneuron in response to a voltage pulse command. (*bottom*) Voltage responses to -50 (blue), 0 (red), and 50pA (purple) current injections, which reveal the characteristic fast spiking of PV+ interneurons. **(d)** Current responses following LSPS glutamate uncaging at the central 8X8 stimulation grid locations surrounding the neuronal soma (indicated by the red square in panel b). Triangle indicates the location of the neuronal soma. (e) Post-hoc validation of parvalbumin expression with immunofluorescence cytochemistry for neurons infused with biocytin during whole-cell recordings. Recorded neurons express GFP (*top*), parvalbumin (*bottom*), and contain biocytin (*middle*). Only PV+ biocytin-labeled neurons were used for analysis.

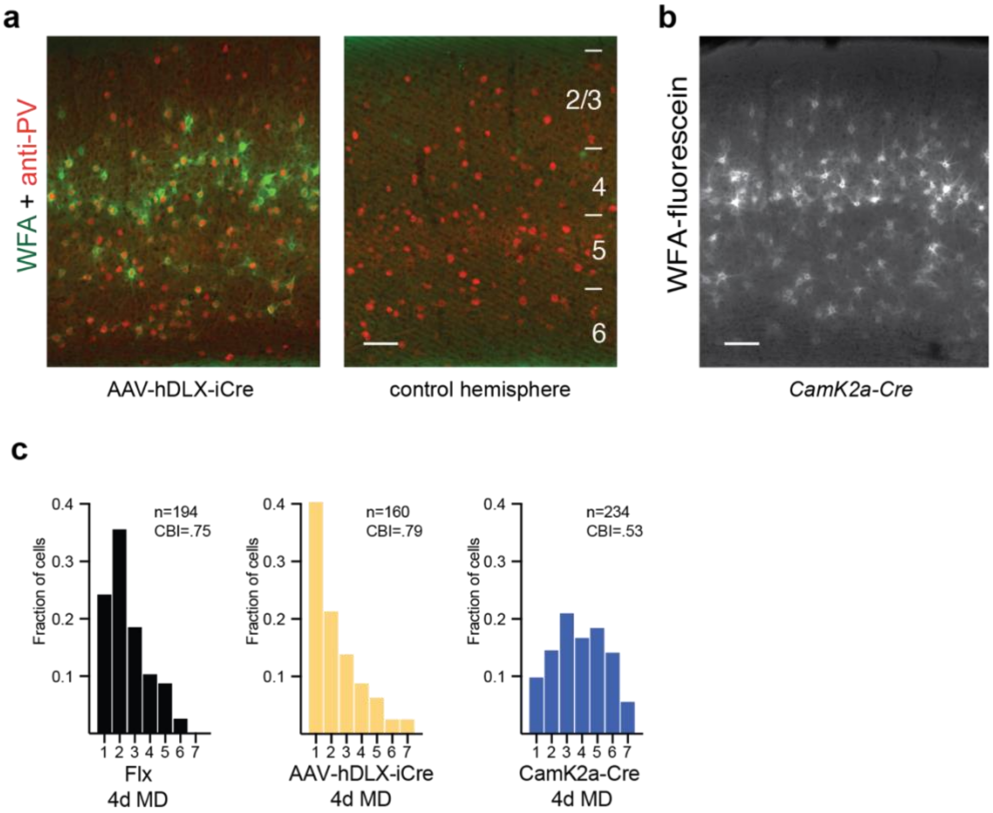


**Figure S3. Deletion of *Acan* in excitatory cortical neurons is sufficient to prevent the closure of the critical period for OD plasticity. (a)** Larger field of view of coronal sections of visual cortex stained with WFA-fluorescein to label PNNs for both hemispheres of mice injected unilaterally with AAVs to express Cre recombinase in inhibitory cortical neurons. Cortical layers are indicated at right. Scale bar = 100 microns **(b)** Larger field of view of a coronal section of visual cortex from an *Acan flx/flx; CamK2a-Cre* mouse stained with WFA-fluorescein to label PNNs. Scale bar = 100 microns. **(c)** OD histograms for the groups of mice receiving 4 days of MD prior to recording that are presented in Figure 3 panel b. CBI is the mean of the number of units (n).
